# Dilated cardiomyopathy remodels cardiac neuronal subtypes in a murine model

**DOI:** 10.1101/2025.08.05.668697

**Authors:** Sachin Sharma

**Author notes:** Corresponding authors: Dr. Sachin Sharma.

## Abstract

The autonomic nervous system is comprised of two parallel chains sympathetic and parasympathetic nervous system that innervates myocardium heterogeneously. However, excessive sympathetic signalling is a hallmark of cardiac dysfunction including myocardial infarction and cardiomyopathies. In current research article, first, we carried out echocardiography on DCM-Tg9 mice to characterize cardiac output parameters and performed scRNAseq analysis in a dilated cardiomyopathy mouse model of heart-failure over wildtype control mice. We analysed differential gene expressions and performed pathway enrichment analysis. The important findings of our research indicate the increment within a specific neuronal subpopulation (NA1b) out of three adrenergic cardiac innervating neuronal subpopulation. We also observed significant increase in M-current potassium ion channel (Kcnq2 ion channel) and genes involved in pathways from oxidative phosphorylation and neurodegeneration pathways. These observations have clearly interpreted that significantly elevated expression of M-current ion channels might be involved with hyperactivated sympathetic signalling in heart-failure mice.

**Highlights:**

1. We identified three adrenergic neuronal subtypes that innervate myocardium and one of these neuronal subtypes, NA1b, predominantly presents during the heart-failure.
2. Our results indicated presence of m-current ion channels (Kcnq2) significantly elevated during heart failure in a murine dilated cardiomyopathy model.
3. The pathway enrichment analyses showed increased expression of genes that are involved in neurodegenerative and oxidative pathways.

## Introduction

Cardiac morbidity and mortality remain one of the largest global burdens of disease according to the data projected in a study^1,2^. Patients with dilated cardiomyopathy (DCM) are prone to develop ventricular tachycardia/fibrillation (VT/VF), a significant cause of heart-failure related deaths^3–5^. After cardiac injury, hyperactive cardiac sympathetic signalling underlies the pathophysiology of adverse LV remodelling and initiation of VT/VF. Currently, available drugs β-blockers and calcium channel blockers attenuate excessive cardiac sympathetic signalling but also impose significant side effects and limited efficacy^5,6^. Therefore, these shortcomings suggest that novel therapeutic strategies that target cardiac-specific neurons rather systemic sympathetic adrenergic signalling blockade have become foundation of the current work.

The intricate role of autonomic nervous system in cardiac pathophysiology has long been established^7–9^. Cardiac-innervating sympathetic neurons that reside in stellate ganglia modules cardiac functions through innervating nerve-fibres^10,11^. Based on their transcriptomic profiling, we identified distinct cardiac-innervating adrenergic neuronal subpopulations in control mice^12^. However, these studies aren’t sufficient to explain whether and how heart-failure remodels cardiac neural subtypes. The current study investigates dynamics of heart-failure induced remodelling of cardiac-projecting neurons in stellate ganglia.

## Methods

### Non-Ischemic Mouse Model of Heart Failure

Animal experiments complied with institutional guidelines and ethical regulations. The study protocol was approved by the UCLA institutional Animal Care and Use Committee. Adult male mice, aged between 8 to 10 weeks, were housed according to the standard laboratory conditions (12h light/dark) with ad libitum access to food and water. Mice strains: C57BL/6J (**000664**) were purchased from Jackson Laboratory and DCM-Tg9 were received from UCLA mice colony stock.

### Transthoracic Echocardiography

Echocardiography was performed on mice using a VisualSonics Vevo2100 system (VisualSonics, Toronto, Ontario, Canada). End Diastolic Diameter (EDD) and ejection fraction (LVEF %) of the left ventricle were assessed to assess left ventricular size and function.

### Retrograde axonal tracing and ganglia dissociation for scRNAseq analysis

We used C57BL/6J and DCM-Tg9 mice strains for tracing the cardiac-innervated neurons in the stellate ganglia and subjected for scRNAseq analysis as per described previously in the “Methods” section^12^.

### scRNAseq analysis of stellate ganglia from DCM vs. WT mice

To map the cell types from our stellate ganglion scRNAseq dataset, we subset the dataset (DCM and WT), and our annotated dataset (WT) to the overlapping highly variable genes^12^. We then used a linear support vector machine (LinearSVM implementation in scikit-learn) with a regularization parameter of 2.0. We trained SVM classifier on our annotated dataset and classified cell types in new dataset. We repeated this process after sub-setting to neurons, to identify the annotated neuronal subtypes in the DCM dataset. The data was then processed as above for dimensionality reduction and visualization (SF1).

To visualize trends in differential expression between DCM and WT neuronal subtypes for ion channels, neuropeptides, and transcription factors, we determined logFC between the average expression of a given gene in DCM vs. WT neuronal subtypes. These logFC values were then visualized with heat-maps. We plotted logFC of the group with the higher expression with respect to the other group. Gene expression in DCM was indicated by blue colour compared to expression in WT as indicated by brown colour. We determined differential gene expression between subtypes in DCM vs. WT mice using the Wilcoxon test (FDR < 0.05). We determined suggestive pathways utilizing suggestive differentially expressed genes (p-value < 0.05) and significant pathways as described above (FDR < 0.05).

### Statistics

Microsoft Excel and GraphPad Prism 9.0.1 were used for data analysis and graph generation. Sample sizes and statistical tests performed are indicated in figure legends. Legend for significance: \**p* ≤ 0.05, \*\**p* ≤ 0.01, \*\*\**p* ≤ 0.001, \*\*\*\**p* ≤0.0001. All data are shown as individual data points.

### Software packages

scRNAseq processing: R, cellranger, Seurat v3. Visualization: python, scanpy, pandas, numpy, matplotlib, seaborn. *Npy* association: python scipy.

## Results

We performed scRNAseq analysis on non-ischemic model of heart-failure (DCM vs. WT) mice using retrograde viral tracing methods.

### 1. Cardiac output parameters

To understand how cardiac SGN subtypes are altered by heart failure, we performed scRNAseq on stellate ganglia from an established transgenic mouse model of chronic non-ischemic heart failure (DCM-Tg9) and control littermates. In this model cardiac dysfunction begins at 7 weeks of age and over, and heart failure at 10 weeks of age, as confirmed by transthoracic echocardiography. We measured cardiac output functions of non-ischemic heart failure mice model (DCM-Tg9)^13^.

Figure **1A** represents (right-side) enlarged chamber size of left ventricle (cardiomegaly) in DCM. We observed progressive decline in ejection fraction and fractional shortening, and increased LV end diastolic diameter and, thinning of left ventricular anterior and posterior walls in DCM compared to wildtype control littermates (Figure **1B**). We also observed progressive fibrosis in heart-failure mice compared to control littermates (Figure **1C**).

**Figure 1.**
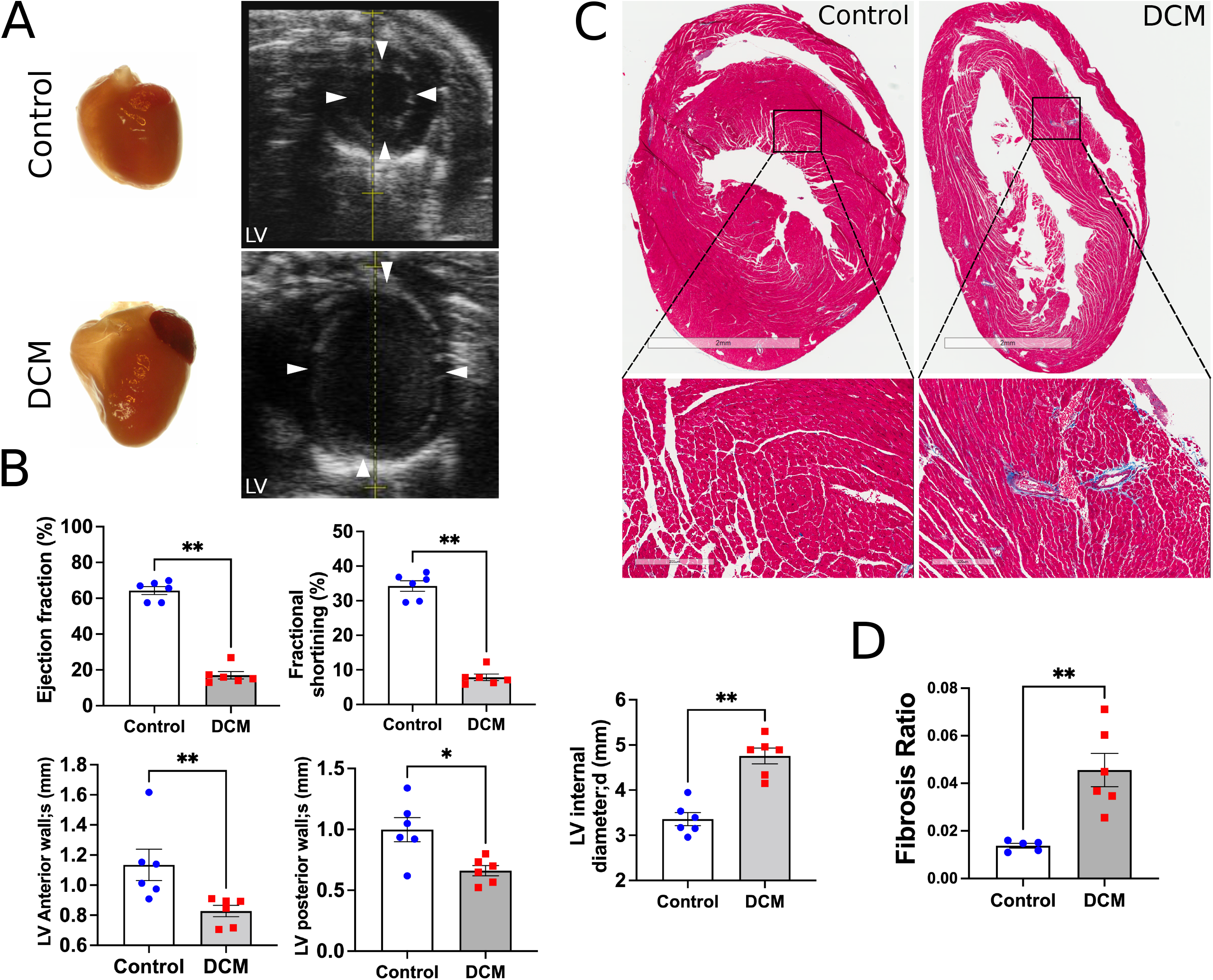
Echocardiography and fibrosis analysis in dilated cardiomyopathy mice (DCM). **A**) DCM hearts grow bigger in size due to increased left ventricular (LV) chamber size. **B**) Ejection fraction, fractional shortening, LV anterior wall, LV posterior wall, and LV internal diameter in wildtype (control) and DCM mice. **C**) Masson’s trichrome staining in heart sections from control and DCM mice. **D**) Fibrosis analysis in control and DCM mice. Data are shown as means ±SEM. For each group 6 mice were used. * p < 0.05, **p < 0.01. Mann-Whitney test was used for the statistical analyses.

### 2. Heart failure differentially impacts cardiac neurons

We performed scRNAseq analysis of stellate ganglia from 10-weeks-old DCM and WT littermates. We identified the same neuronal subtypes from our prior analysis with a support vector machine (SVM) classifier as described previously in Methods section^12^ (Figure **2A**). However, we observed that unlike control mice where cardiac subtypes are equally distributed among NA1a, NA1b, and NA3 subtypes, the NA1b subtype become predominant in DCM mice (Figure **2B**). We found that NA1b neurons represents a greater proportion of DCM cardiac neurons, while NA1a and NA3 subpopulations were reduced in stellate ganglia. Since NPY has an important role in heart failure, we characterized the distribution of NPY expression in tertiles (low, medium, and high) across the cardiac subtypes in DCM mice compared to WT animals. While most NA1a and NA1b neurons remained high and low NPY expressing, respectively, most NA3 neurons were high NPY expressing, suggesting remodelling of this subtype of neurons towards greater NPY expression (Figure **2C**).

**Figure 2.**
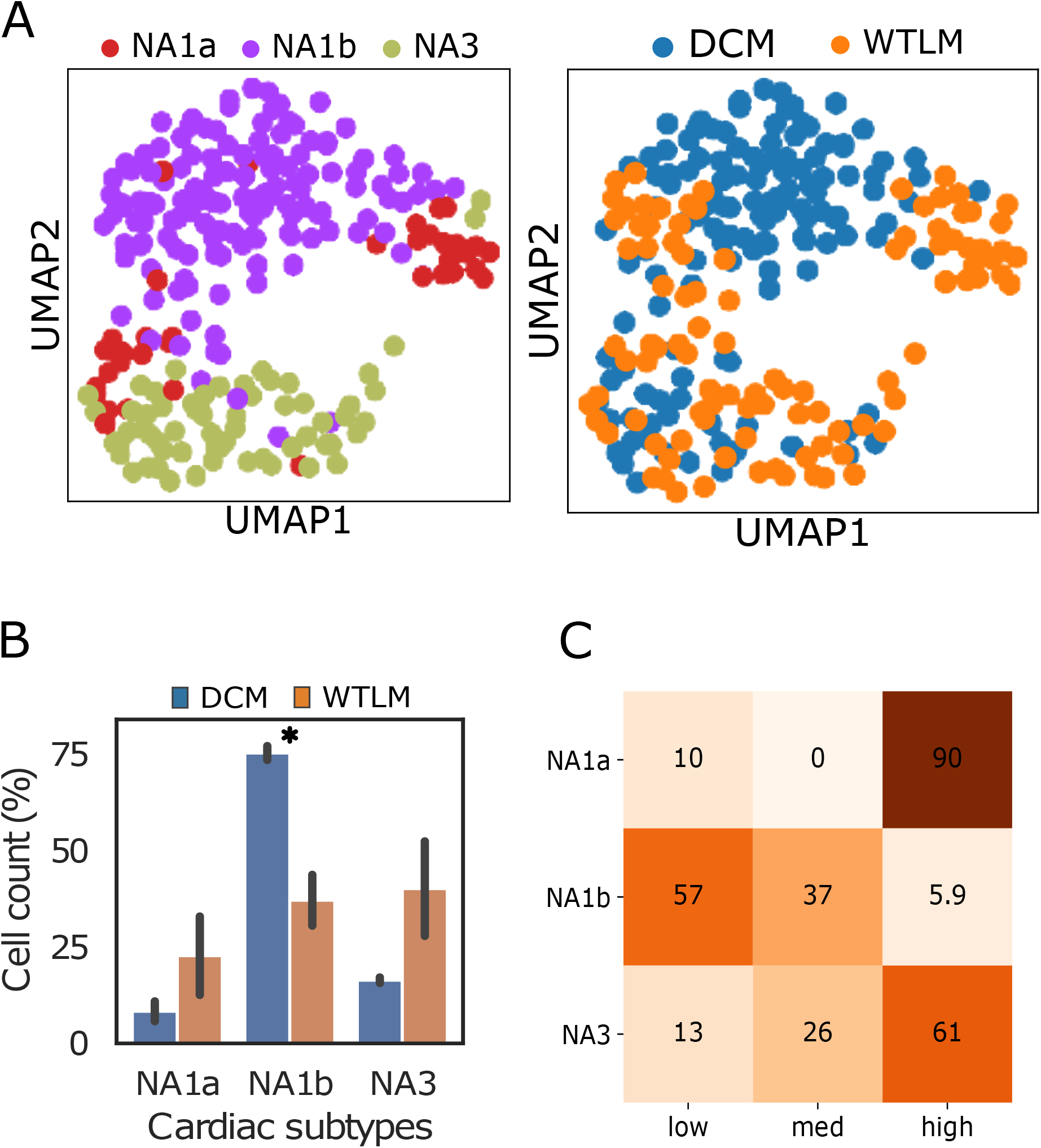
Cardiac neuronal subtypes and their transcriptomic profile in the stellate ganglia from dilated cardiomyopathy (DCM) mouse. **A**) UMAP plot shows the cardiac subpopulations in DCM and WT mice. In DCM mice, NA1b becomes the predominant neuronal subtypes. **B**) Proportion of neuronal subtypes cell count in DCM and WT littermates. **C**) Distribution of cardiac subtypes across the quantiles of NPY expression in DCM mice.

### 3. Gene enrichment and Pathway analysis

We analysed a subset of differentially expressed genes (DEGs) in cardiac SGNs from DCM and WT animals. We found significant DEGs only in the NA1b subtype with the top 10 upregulated and downregulated genes in NA1b subtype in DCM animals shown in Figure **3A, B**.

Our data suggest that lack of DEGs in other cardiac subtypes (NA1a, NA3) is likely due to relatively low cell numbers of these subtypes in WT and DCM animals. Interestingly, we observed a specific and significant increase in the expression of M-current conducting ion channel Kcnq2 in DCM mice (Figure **3C**). Along with Kcnq2, a slowly activating and deactivating potassium channel that plays a critical role in the regulation of neuronal excitability and synaptic transmission across the chemical synapses, we found observed trends indicating increased expression of voltage gated sodium channels (Nav1.1, Nav1.2), potassium channel (Kv11.3), calcium channel (Cav2.2), and vesicle transport proteins (Ap2a2, Snap25) (Figure **3C**). These alterations in ion channel expression in this subtype implicate the NA1b subtype in the alterations of sympathetic neuronal excitability demonstrated in the setting of heart failure.

**Figure 3.**
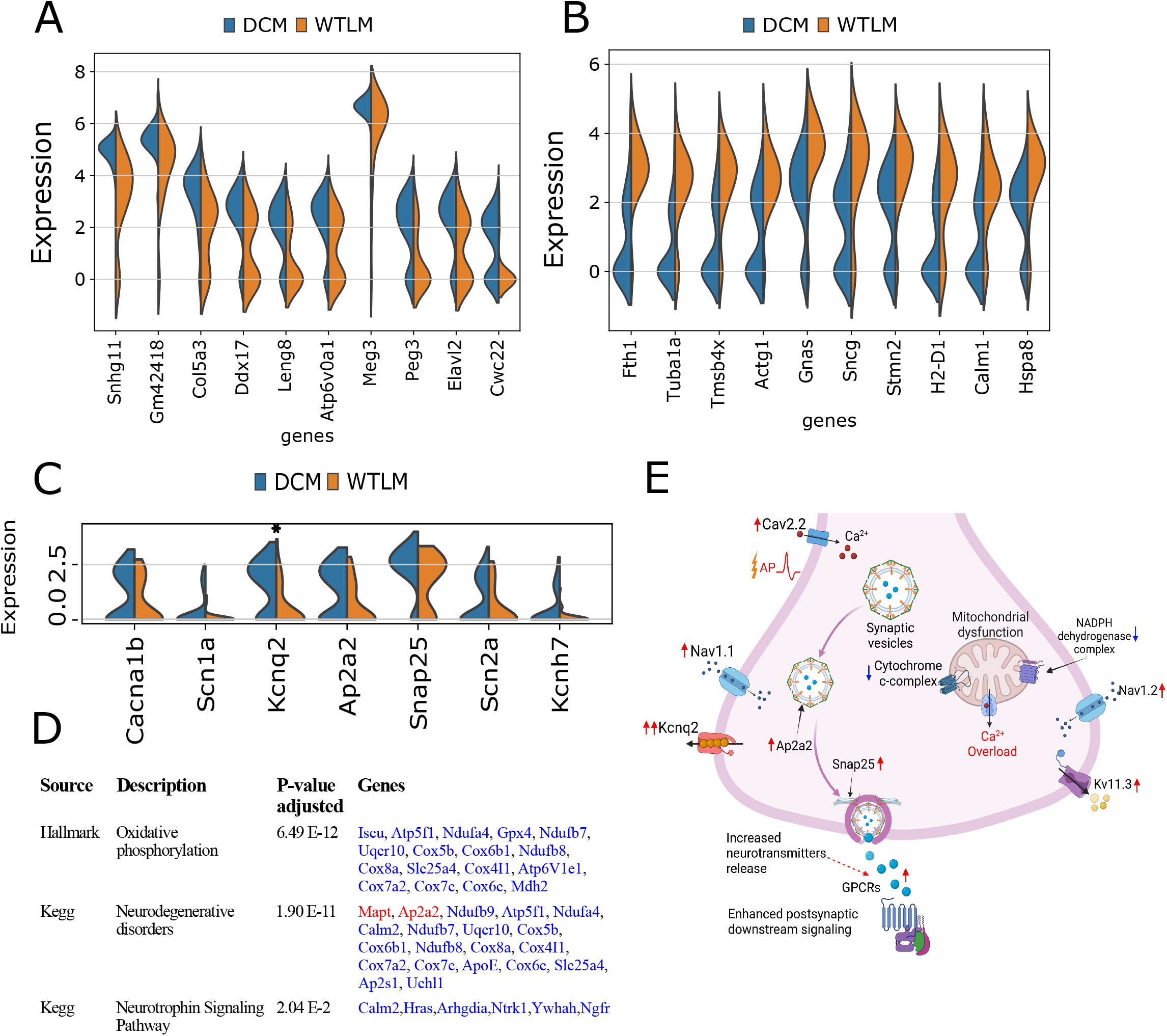
A: B, Violin plots show top ten significantly upregulated genes (**A**) and downregulated genes (**B**) in NA1b subtypes in heart failure. **C**) Violin plot indicating differential expression of ion channels and vesicular transport genes in NA1b subtype in heart failure. **D**) Relevant significantly enriched pathways in NA1b subtype. **E**) A schematic diagram of the potential mechanism underlying the increased neuronal activity in NA1b subtype, elucidated based on the DEGs and pathway analysis in heart failure mice. Data are shown as means ±SD. * p < 0.05 compared to control. (WT: n=4 and DCM: n=2)

**Figure 4:**
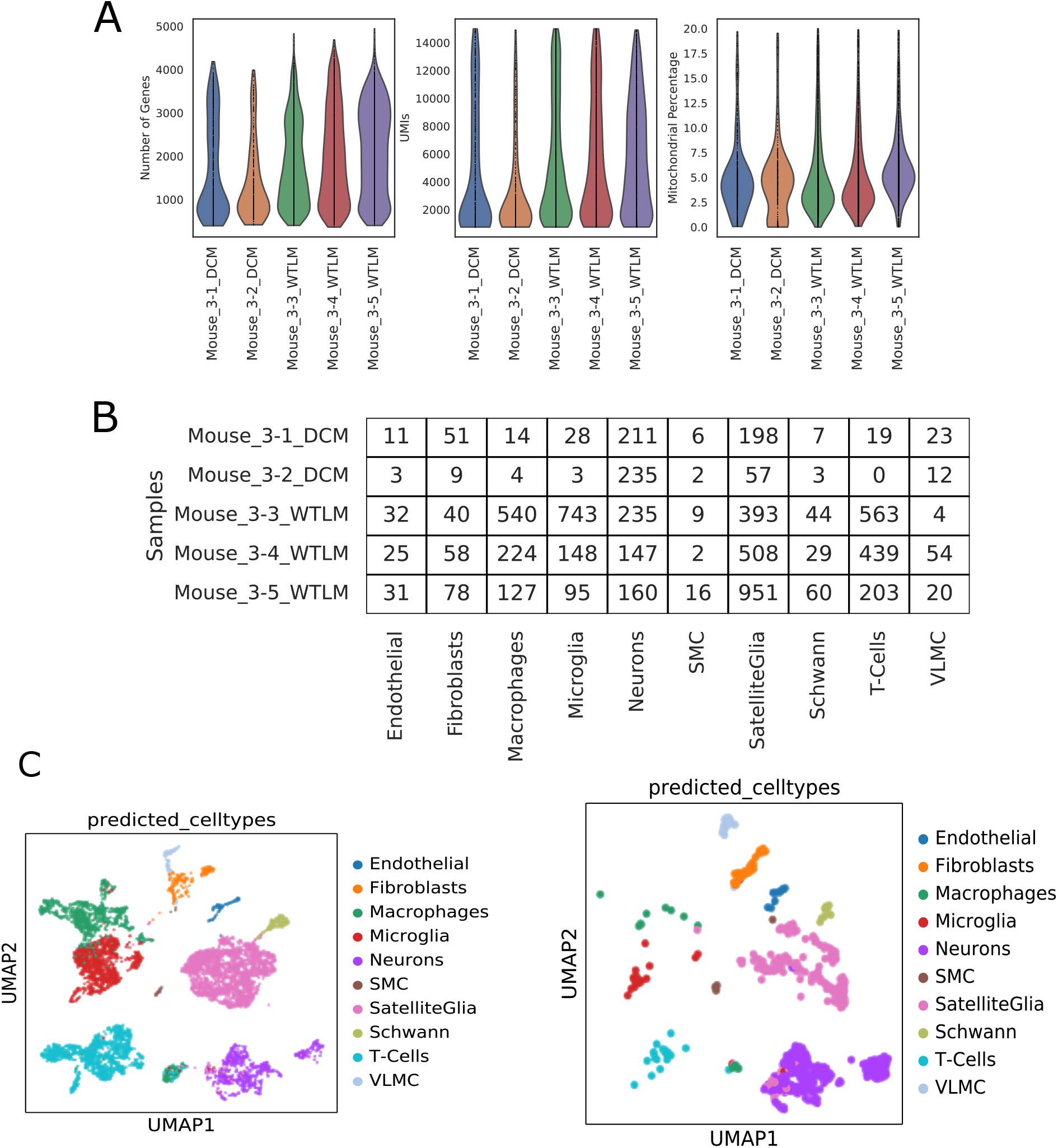
A schematic diagram showing representation of workflow that has been carried out in this manuscript.

We performed pathway enrichment analysis on the suggestive DEGs (p < 0.05) stellate ganglion neurons from DCM mice and found enriched pathways (FDR < 0.05) in only NA1a and NA1b subtypes. We identified relevant significantly enriched pathways from NA1b subtypes where three or more than three genes were affected. We selected over-representative pathways: oxidative phosphorylation (FDR=6.49e-12), neurodegenerative disease (FDR=1.90e-11), and neurotrophins signalling (FDR=2.04e-2) pathways from NA1b subtype (Figure **3D**). In these pathway enrichment studies, we found significant downregulation of mitochondrial genes in cardiac neuronal subtypes (NA1a, NA1b, NA3), suggesting an impairment in the metabolic function during the heart failure (Figure **3D**). Based on the DEGs and pathways enrichment analysis, our studies suggest that perturbed excitability of NA1b sympathetic neurons may enhance release of neurotransmitters that amplify chronic sympathetic signalling in heart failure.

## Discussion

The neural regulation of myocardium is compartmentalized in layers spanning from intrinsic cardiac nervous system (located within the pericardium), extra-cardiac ganglia, and central nervous system (brain and spinal cord)^14,15^. The postganglionic sympathetic neurons residing within stellate ganglia receive pre-synaptic inputs from preganglionic sympathetic fibres originating from spinal cord and transmute post-synaptic signals to target innervating organs^16^. Postganglionic sympathetic neurons residing within the stellate ganglion innervate predominantly myocardium and other tissue-beds in visceral organs as revealed by studies carried out using retrograde tracers^11,12^.

As such, we investigate how heart failure influences the distribution of cardiac SGN subtypes, and how NPY expression is altered in NPY high and NPY low/negative subsets. Two main findings from our study provide insights into potential mechanisms driving elevated NPY levels in the setting of chronic cardiac injury. First, we find that the 3 cardiac SGN subtypes (NA1a, NA1b, and NA3) remain preserved, however, heart failure shifts the relative distribution across the 3 subtypes towards one dominant one (NA1b). Importantly, NA1b, which had low levels of NPY expression in control littermates, exhibited increased NPY expression albeit to levels lower than that seen in NA1a. Taken together, our findings suggest that the transcriptomes of cardiac SGNs generally shift to amplify the subtypes and numbers of cells that express NPY. Given the profound actions of NPY as a potentiator of adrenergic signalling, our findings provide a basis for the prognostic role of elevated circulating or coronary sinus levels of NPY in identifying patients with severe sympathoexcitation for whom outcomes are dismal. Our findings are in line with those of Davis et al^17^, who performed scRNAseq in rats with spontaneous hypertension (SHR) and Wistar controls. In this study, cardiovascular SGNs were not specifically identified for characterization, however, SGNs in the SHR model exhibited functional (increased excitability and tonic firing) and transcriptomic alterations (reduced transcript levels of genes encoding the M-current, subunits KCNQ2, KCNQ3, and KCNQ5). In our study, we found increased expression of KCNQ2 in heart failure animals, contrary to decreased expression observed in the SHR model. This suggests that although sympathetic activation is characteristics of both hypertension and chronic heart failure, the mechanisms, at the neuronal level, that drive sympathetic activation may differ fundamentally at the neuronal level.

Our findings of a dominant neuronal subtype with greater NPY expression in the setting of heart failure may have significant clinical implications. In patients with chronic heart failure or arrhythmias, systemic pharmacologic blockade of neurohormonal activation and sympathetic signalling, is a mainstay of treatment^18–20^. While these drugs have proven lifesaving benefits, they result in substantial side effects such as hypotension, mental incapacitation, fatigue, weakness, and sexual side effects to name a few^21^. Of note, therapies that target the stellate ganglion, rather than systemic blockade, have shown significant benefit and are increasingly used clinically. Yet, these approaches also target the entire stellate ganglion with potential off target issues to the other tissue beds and organs innervated by neurons in the stellate ganglion. Hence, more specific targeting of neurons that innervate the heart, and drive the excessive chronic sympathetic signalling characteristic of chronic cardiac injury are needed. Our study presents one such candidate, the cardiac SGN subtype NA1b, which becomes the dominant cardiac neuronal subtype in heart failure. Analysis of the transcriptome of this cell identifies potential targets that can be modulated directly or used to identify this cell type for other forms of intervention, for example, biologic agents. Coupling such cell specific targeting with localized delivery or such agents to the stellate ganglion via minimally invasive approach (for example as used with stellate ganglion block) provides novel avenues to target sympathetic excess in chronic cardiac injury.

The present study has limitations. First, changes in transcript levels in DCM vs control stellate ganglia identified by scRNAseq do not connote functional changes, hence our findings imply but do not prove functional differences in the setting of heart failure. However, the extensive prior published literature showing functional changes in SGNs following cardiac injury support functional consequences of the differences in gene expression we identified in this study. Second, although we find that the NA1b subtype becomes dominant in the setting of heart failure, we postulate but do not have direct evidence that they are responsible for chronic hyperactive cardiac sympathetic tone.

## Conclusion

In summary, we identified, that cardiac neuronal subtypes in the SGN are differentially impacted by heart failure, yielding a dominant cardiac SGN subtype that may represent a novel target for adrenergic blockade in heart failure. These findings expand our understanding of cardiac sympathetic control mechanisms and suggest that cell-specific targeting of sympathetic neurons may offer new therapies for heart failure.

## Author acknowledgement

SS has conceptualized the study, edited and drafted the final version of the manuscript.

## Funding statement

The current research has been funded by a specific grant received from American Heart Association (Grant ID: 906065) to Sachin Sharma during his postdoctoral work at UCLA Cardiac Arrhythmia Centre, Los Angeles, 90055, USA.

---

**SF1:**
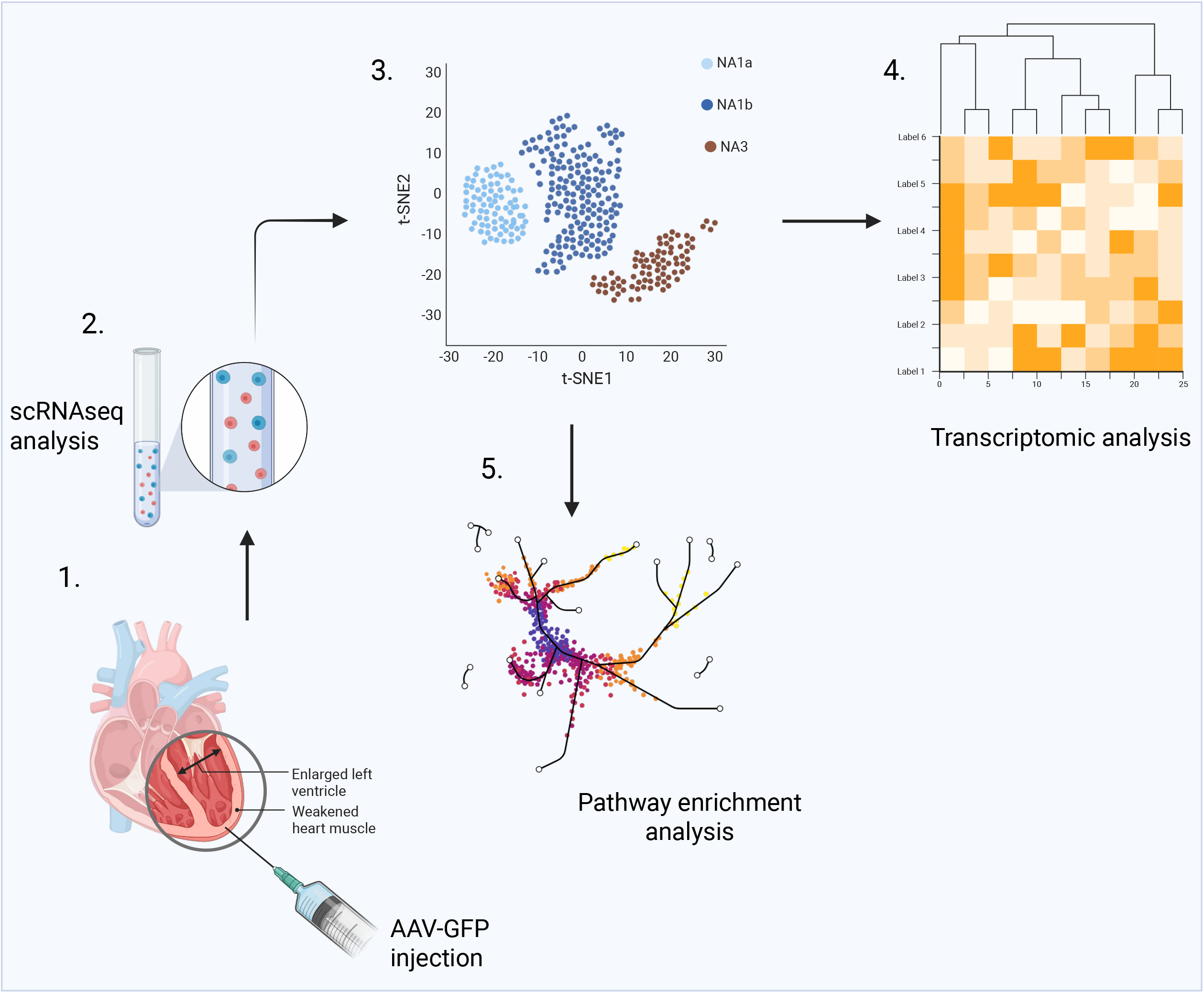
scRNAseq quality control. Samples are indicated with their batch, sample number, and disease status. **A**) Violin plots showing the distribution of the number of genes, UMIs, and mitochondrial percentage per cell in the experiment with WT mice (n=4) and in DCM mice (n=2).

## Notes

### Competing Interest Statement

The authors have declared no competing interest.

